# Generation and Assessment of High-Quality Fat-Tailed Dunnart Oocytes Following Superovulation in Prepubertal Animals

**DOI:** 10.1101/2025.03.11.642722

**Authors:** Jun Liu, Namdori Mtango, Emily L. Scicluna, Sara Ord, Andrew J. Pask

## Abstract

The fat-tailed dunnart, *Sminthopsis crassicaudata*, is a mouse-sized, polyovular, solitary dasyurid marsupial found in central and southern Australia. With the establishment of a chromosome-scale genome assembly, induced pluripotent stem cells, and targeted genetic editing, the dunnart is emerging as the laboratory marsupial model for comparative developmental, reproductive and conservation biology. The development of assisted reproductive technologies (ART) are critical to achieving these goals in this species. ART requires a large number of mature oocytes which are typically collected through stimulated and synchronised female reproductive cycles. While protocols for induced-ovulation or superovulation are standard in many placental mammals, there are no methods to date designed for marsupials. In the present study, prepubertal dunnarts were stimulated with pregnant mare serum gonadotrophin and human chronic gonadotrophin across 6 different treatment regimens. Our best regimen resulted in over 70% of prepubertal dunnarts ovulating with 82% normal oocytes. When the primed females were mated with stud males, 4-cell stage embryos were collected 48 h post-hCG administration. At around 96 h post-hCG, 50% (n=8) and 78% (n=9) of the embryos developed to blastocysts. Our results demonstrated successful stimulation of ovulation and mature oocyte collection in prepubertal dunnarts. Furthermore, we confirmed developmental competence of the induced ovulated oocytes through to at least the blastocyst stage. These findings represent the first robust hormonal regimen for predictable oocyte generation in any marsupial and will significantly contribute to the use of the dunnart in developmental and conservation biology.

## Introduction

The fat-tailed dunnart (*Sminthopsis crassicaudata*, hereafter referred to as the dunnart) is a mouse-sized, polyovular, solitary dasyurid marsupial found in central and southern Australia. Female dunnarts reach sexual maturity at approximately three months of age and have a prolonged estrus cycle of 31 days [1–4], compared to four to five days in the mouse [5]. This species is an excellent candidate for laboratory-based marsupial research animal, owing to its simple and robust husbandry, unique gamete biology [6–8] and biological features such as distinctive mode of reproduction, developmental heterochrony in craniofacial and limb patterning, and altricial stage at birth [9–12]. The dunnart is further supported by additional resources, including chromosome-scale genome assembly [13], induced pluripotent stem cells [14–16], and targeted genetic editing [17]. Thus, the dunnart is emerging as a robust marsupial model for comparative developmental and conservation biology studies.

Harvesting sufficient oocytes is a critical step for the implementation of assisted reproductive technologies (ARTs), including in vitro fertilization (IVF), intracytoplasmic sperm injection (ICSI), cloning by somatic cell nuclear transfer (SCNT) as well as embryo injection of CRISPR reagents for the generation of gene targeted animals. Previous attempts were made to control the female reproductive cycle as a source of oocytes and embryos in adult dunnarts. This includes the removal of young from the mothers’ pouch (removal of pouch young; RPY) which suspends lactation and induces follicular growth, reactivating the natural cycling in polyestrous marsupials [18, 19]. RPY was also used to synchronize cycling between females [20]. Although long established as a research tool, the accuracy of the subsequent ovulation and mating following RPY are imprecise and occur over a range of several days in most species [18]. In a recent study, body weight fluctuations could be used as an ovulation indicator and achieved >75% success rate for embryo collections [12]. The body weight fluctuation pattern across an entire reproductive cycle consists of a temporary weight drop indicating ovulation, progressive weight increases during pregnancy and a sharp weight dip at birth. The daily body weight monitoring is time-consuming and has a significant cumulated burden for the animals exacerbated by the protracted estrus cycle in the dunnarts (around 31 days). In addition, weight fluctuations are mostly only consistent when female dunnarts are caged with a male, and therefore for embryo collections, but this is not ideal for unfertilised oocyte collections in support of advanced ART. Another disadvantage with this method is that body weight changes are impacted by other factors, such as husbandry conditions and frequent animal handling which limits its accuracy as a practical tool for detecting reproductive status.

Methods for collecting oocytes in placental mammals such as rodents [5, 21, 22], rabbits [23, 24] and livestock [25–27] are well established. In mice, the most prevailing induced ovulation protocol involves injection of prepubertal females with pregnant mare serum gonadotropin (PMSG) to stimulate follicular development, followed by an injection of human chorionic gonadotropin (hCG) to trigger ovulation [21]. This results in the supraphysiological generation of oocytes (superovulation) for use in subsequent workflows. Prepubertal mice are used as they do not exhibit the regular estrus cycles seen in adult females. This lack of endogenous hormonal cycling ensures there is no natural secretion of gonadotropin (FSH and LH) that could conflict with the exogenous hormones administered for superovulation, leading to more consistent and reproducible results. There have been attempts to induce ovulation using exogenous gonadotrophins in marsupials, namely in adult fat-tailed dunnarts [28–30] and stripe-faced dunnarts [31–33]. However, a robust hormone regimen for induced ovulation/superovulation has not yet been established as a reliable means for the control of estrus in dunnarts. Significant research with a focus on dose refinement and hormone priming timing is required. In addition, the development of an estrus synchronization strategy in adult animals is a prerequisite prior to gonadotrophin stimulation. In this way, the PMSG administrated for superovulation would be in sync with the endogenous hormone environment to avoid conflicting signals for the ovaries. Gonadotrophin releasing hormone (GnRH) analogue-based female synchronization technologies for marsupials require further investigations [34, 35], since they would provide a significant advance for dunnart ART development.

The fat-tailed dunnart has been a focal species for advancing assisted reproductive technologies and developing next generation conservation strategies for marsupials. These advancements are crucial for preserving and restoring genetic diversity, which could support the conservation of this species and other threatened marsupials. However, challenges remain in obtaining large numbers of mature oocytes from fat-tailed dunnarts, particularly due to variability in ovarian responses to stimulation and the resulting quantity and quality of the oocytes. Despite the widespread use of superovulation in prepubertal mice, this approach has previously not been attempted in prepubertal dunnarts. This study aimed to develop a superovulation protocol to stimulate complete oocyte development in prepubertal fat-tailed dunnarts. The developmental potential of the oocytes obtained was assessed based on their maturation, fertilization competence, and ability to support early-stage embryogenesis.

## Materials and Methods

### Animals and Experimental Groups

Female fat-tailed dunnarts, housed and bred in the Biosciences 4 animal facility at the University of Melbourne, were used throughout the study. The animal rooms were maintained at a temperature of 25 °C and humidity of 50% under artificial lighting (16 h light (5:00 am to 9:00 pm) and 8 h dark (9:00 pm to 5:00 am)). Ethics approval for the study was obtained from the Animal Research Ethical Committee of the University of Melbourne (Project ID 26863 and 26864). Prepuberal dunnarts aged between 70-85 days were primed with 1 IU PMSG (ProSpec, Ness Ziona, Israel) for 4, 6, or 8 days either through a single injection (given intraperitoneally; IP) or IP injections every two days. This was followed by IP administering 1 IU hCG (ProSpec) on day 4, 6 or 8, respectively. Oocytes were collected from oviducts and/or uteri 10-14 hours post hCG administration. For embryo collection, the superovulated females were caged with stud males and embryos were collected 48 hours after hCG administration.

### Oocyte retrieval

Oocytes were manually flushed out of the uteri and oviducts using 30-½ gauge needles with M2 medium (Sigma). After washing 3 times in M2 medium, the cleaned oocytes were allocated to 20 μL droplets of medium and covered with mineral oil in 35mm dishes (#3001; Falcon). The numbers of ovulated oocytes were counted and classified based on their morphology; oocytes with irregular shape, dark cytoplasm or fragmentations were classified as abnormal. Images of oocytes were captured using an inverted microscope (Olympus X73). The oocyte diameter was measured using FIJI software [36]. The mean diameter was estimated by taking of the average from two perpendicular diameters measured for each oocyte.

### Spindle fluorescence staining

Oocytes were fixed in 4% paraformaldehyde (Sigma) in phosphate buffered saline (PBS, pH 7.4) overnight at 4 °C and permeabilized in 0.5% Triton X-100 (Sigma) for 20 min at room temperature. Oocytes then were blocked in PBS supplemented with 1% BSA, 0.1% Tween 20 (Sigma) and 0.01% Triton X-100 (PBST) for 1h and incubated with anti-α-tubulin-FITC antibody (Sigma, 1:200 dilution) at 4 °C overnight. After washing in PBST, the oocytes were counterstained with 10 µg/ml Hoechst 33342 (Sigma) for 10 min. Finally, the oocytes were transferred to a µ-Slide 8 Well (Ibidi) and observed under a laser scanning confocal microscope (Nikon).

### Embryo retrieval and embryo culture

Embryos were released from the uteri by opening the uteri with scissors. After washing 3 times in M2 medium, the cleaned embryos were incubated in 30 μL drops of modified potassium simplex optimized medium (KSOM) [37] supplemented with 0.4% (w/v) fatty-acid free BSA (MP Biomedicals), 0.5× of NEAA and 0.5× of EAA (ThermoFisher) for dunnart embryo culture (designated dKSOM1/dKSOM2 sequential medium, Supplemental Table S1) covered with mineral oil in 35mm Falcon culture dishes (#3001) at 35°C in a 5% CO_2_:6% O_2_ atmosphere.

### Vaginal lavage cell staining

Females were gently picked up and 20 μL of PBS was expelled at the opening of the urogenital canal using a pipette. The fluid was withdrawn back into the tip and the process was repeated 2-3 times. The lavage was then placed on a glass slide and allowed to completely dry at room temperature. After drying, the samples were stained in 0.1% crystal violet (Sigma) for 3 min, washed with water for 1 min, and then mounted with glycerol and coverslip. The lavage samples were examined under a light microscopy (Motic) to determine the relative abundance of nucleated epithelial cells, cornified epithelial cells (CEC) and leukocytes.

### Histology

Ovaries obtained from the superovulated dunnarts were fixed in Bouin’s solution (Sigma) for 24 h, washed in 70% ethanol several times and then embedded in paraffin, serially sectioned at 5 μm, and stained with hematoxylin and eosin. The sections were scanned and examined for the presence of large antral follicles (AF) and corpus luteal (CL) using the Pannoramic Scan II (3D Histech) at UoM Histopathology and Slide Scanning Service and the free viewing software SlideViewer (3D Histech).

## Results

### Oocyte yield and maturity

A total of 44 prepubertal female dunnarts aged between 70-85 days post-partum were injected with one of 6 regimens of 1 IU of PMSG (Fig. 1): a single injection of PMSG followed by hCG stimulation after 4 days (abbreviated as 1P4, n=2), hCG stimulation after 6 days (1P6, n=2), and hCG stimulation after 8 days (1P8, n=2); or PMSG injections every-two-days for 4 days (2P4, n=5), 6 days (3P6, n=19) and 8 days (4P8, n=9) before hCG injection. In all cases oocytes were collected 10-14 h post-hCG injection.

**Figure 1.**
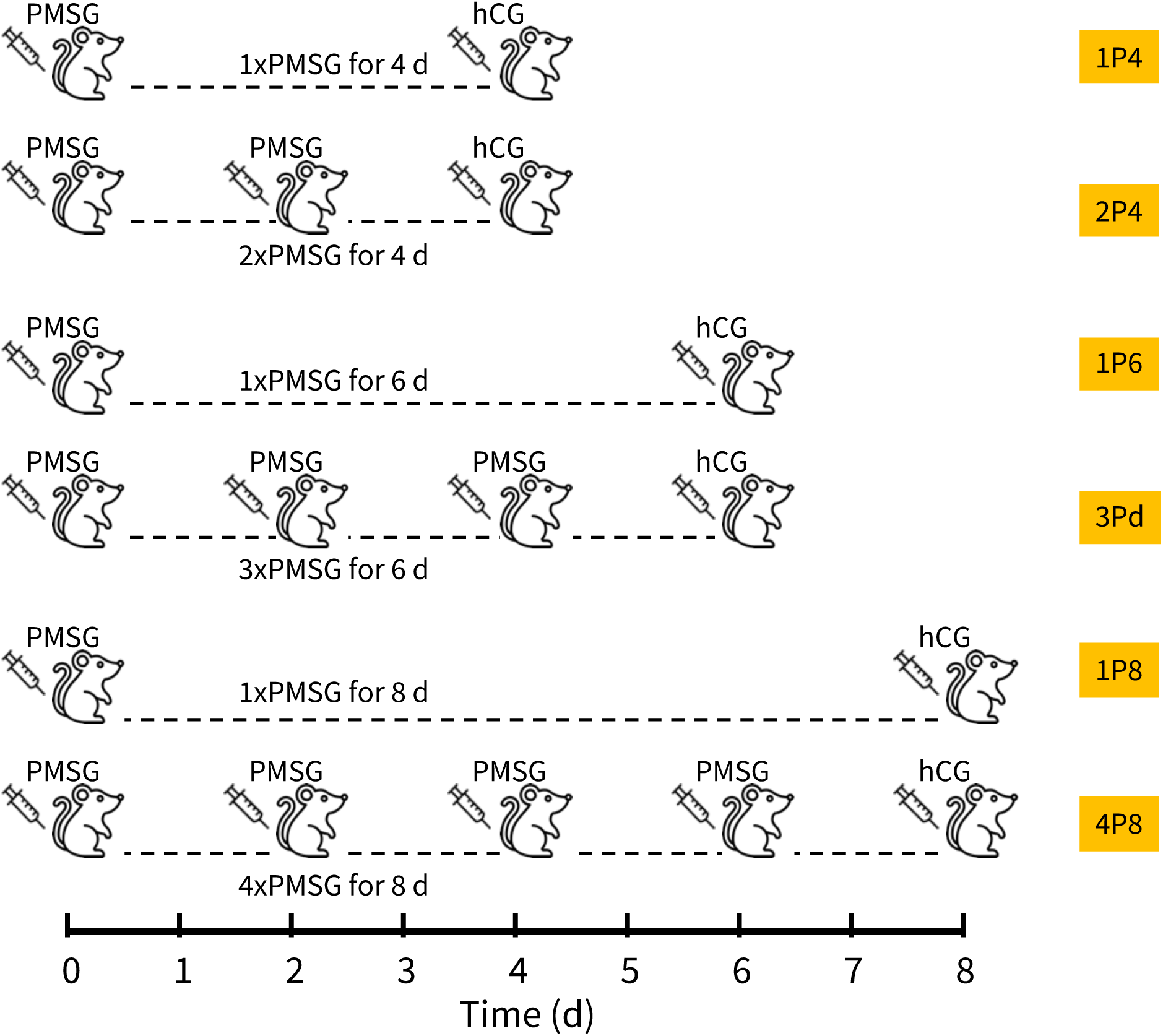
A timeline scheme for hormonal priming: Female dunnarts were primed with 1 IU PMSG in either a single injection on day 0, or once every two days from day 0 followed by a single hCG stimulation 4, 6 or 8 days later.

Among the six groups examined, 8-day PMSG priming regimens (1P8 and 4P8) gave the best results in terms of the percentage of female dunnarts that ovulated (88.9-100%) and the numbers of collected oocytes (mean 13-18 oocytes per dunnart), followed by 6-day priming regimens (1P6 and 3P6) with 50-73.7% ovulation rate and mean 9-13 oocytes collected per dunnart, and 4-day priming regimens (1P2 and 2P2) with 0-20% ovulation rate and mean 0-0.4 oocytes collected per dunnart (Table 1, Fig. 2-A). These data suggest that follicle development was largely not completed in 4 days under the PMSG stimulation, and therefore almost no oocytes were collected. After 6-day and 8-day PMSG priming, 82-100% and 34% of the oocytes displayed normal morphology, respectively (Table 1, Fig. 2-B, C). This indicated that the proportion of oocytes with normal morphology was affected by the PMSG priming duration. Thus, the 6-day PMSG priming protocol resulted in highest rate of production of normal oocytes than the 8- and 4-day priming protocols. There were no significant differences between the single PMSG injection (n=2) and injections every-2-day across the same priming periods (n=19).

**Figure 2.**
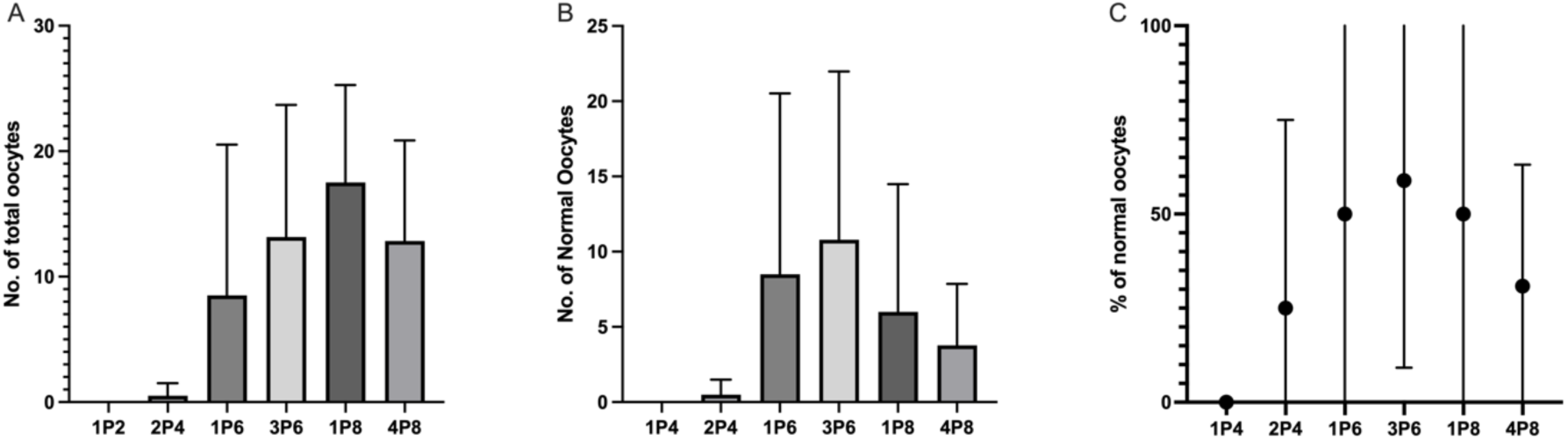
Results of superovulation for each treatment in the prepubertal dunnarts. A) Mean number of total oocytes retrieved per dunnart. B) Mean number of normal oocytes collected per dunnart. C) Percentage of oocytes with normal morphology.

**Table 1.** Results of superovula.on by each treatment in the prepubertal dunnarts.

| Experimental group | No. of females<br>ovulated/tested (%) | Mean ± SD no. of oocytes | | | |
| --- | --- | --- | --- | --- | --- |
|  |  | Total | Normal | Abnormal | % of Normality |
| 1P4 | 0/2 (0) | 0 ± 0 | 0 ± 0 | 0 ± 0 | 0 |
| 2P4 | 1/5 (20) | 0.4 ± 0.9 | 0.4 ± 0.9 | 0 ± 0 | 100 |
| 1P6 | 1/2 (50) | 9 ± 12 | 9 ± 12 | 0 ± 0 | 100 |
| 3P6 | 14/19 (73.7) | 13 ± 11 | 11 ± 11 | 2 ± 6 | 82 |
| 1P8 | 2/2 (100) | 18 ± 8 | 6 ± 9 | 12 ± 16 | 34 |
| 4P8 | 8/9 (88.9) | 13 ± 10 | 4 ± 5 | 8 ± 11 | 34 |

Dunnart oocytes were collected by flushing the oviducts and uteri with M2 medium. A total of 2, 17, 12, 205, and 49 normal oocytes were retrieved from the 2P4, 1P6, 1P8, 3P6, and 4P8 treatments, respectively (Supplemental Table S2). The mean diameter of the dunnart oocytes was 172.2 ± 3.7 µm (mean ± SD, n = 98) with a range of 164.3-180.5 µm in 3P6 group; and 157.1 ± 15.6 µm (n = 29) with a range of 133.6-183.1 µm in 4P8 treatment group (Supplemental Table S2). Dunnart oocytes retrieved from all treatment groups share similar morphological features (Fig. 3A). The dunnart oocyte is surrounded by two extracellular coats: a zona pellucida and a mucoid coat. A large membrane-bound vesicle, named the deutoplast [38, 39] was present in the ooplasm. The first polar bodies can be seen in the thin perivitelline space of oocytes, indicating the completion of the first meiosis and reaching MII stage. Furthermore, barrel-like spindle arrangements indicative of metaphase with well-aligned chromosomes were present in most oocytes (Fig. 3B and C) imaged with immunofluorescent staining and confocal imaging.

**Figure 3.**
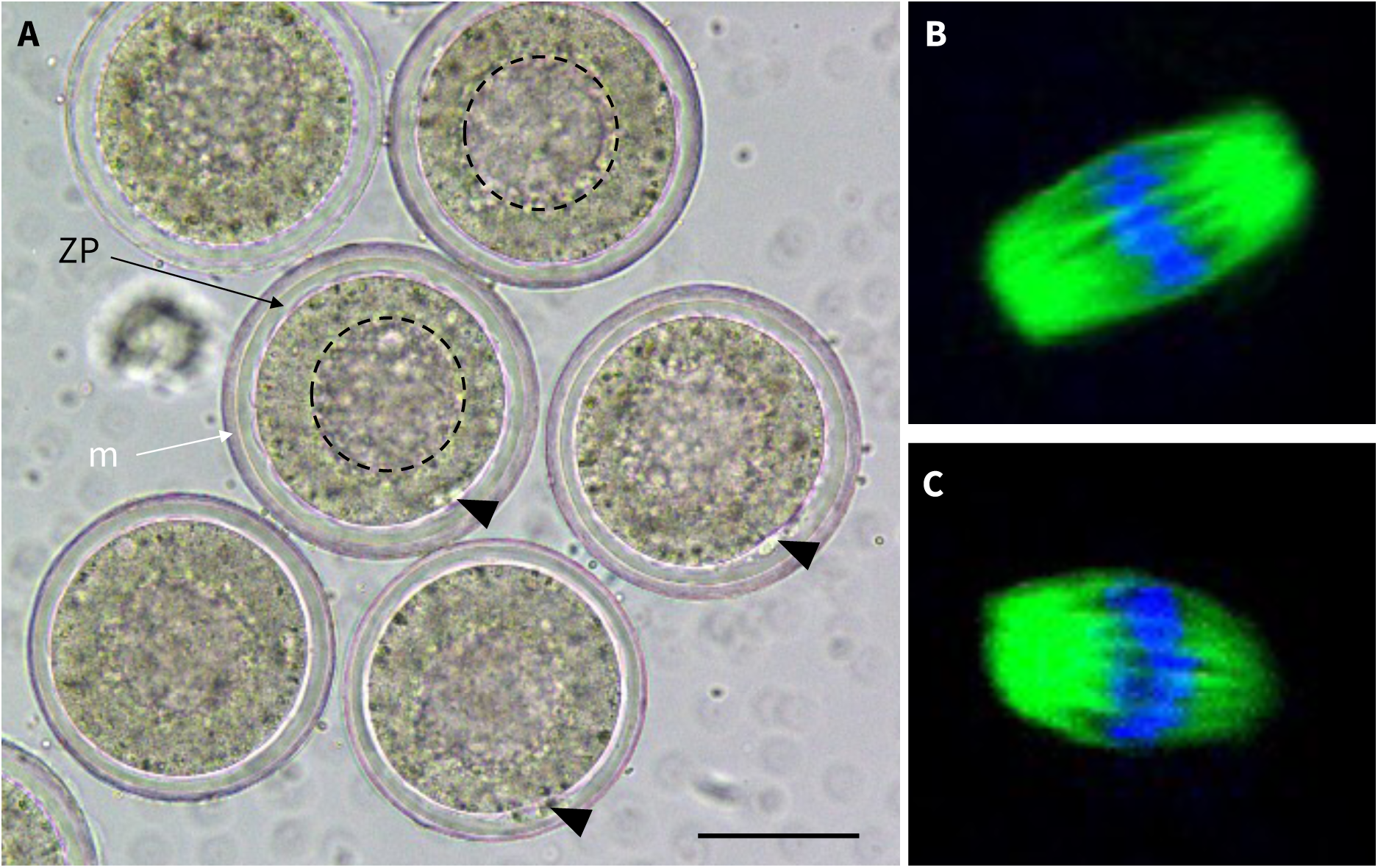
The dunnart oocytes retrieved from prepubertal animals after superovulation. A) Metaphase II oocytes with the first polar body (arrowheads) in the thin perivitelline space, large deutoplasm vesicle (dashed circles) in ooplasm, surrounded by zona pellucida (ZP, back arrow) and mucoid coat (m, white arrow) (bar = 100 µm). B) and C) Representative images of spindle morphology and metaphase chromosome alignment in oocytes at the MII stage. Oocytes were immunostained with the α-tubulin-FITC antibody (green) to display the spindles and counterstained with Hoechst 33342 (blue) to visualise the DNA/chromosomes.

### In vivo fertilization and embryo culture

To assess the capacity of the mature oocytes produced from prepubertal female dunnarts after superovulation in the optimal 3P6 treatment group, females were mated with stud male dunnarts in a 1:1 ratio. The 4-cell stage embryos were collected 48 h post-hCG administration and cultured in dK1 medium (Fig. 4-A). The dunnart embryos developed to 8-cell stage (Fig. 4-B) 20 h later and were subsequently transferred to dK2 medium for blastocyst formation (Fig. 4-C and D) at ∼96 h post-hCG injection. The results are summarised in Table 2. A total of 43, 4-cell embryos were retrieved from two females (with 23 and 20 embryos respectively) in two replicates. Seventeen of the 4-cell embryos were cultured in dK1 medium (the remaining 26, 4-cell embryos were cultured in other media, data not shown). All 4-cell embryos in dKSOM1 medium cleaved to 8-cell or16-cell embryos by the next day. The embryos were then transferred to dKSOM2 medium for blastocyst formation. At approximately 96 h post-hCG administration, 11 of 17 (64.7%) embryos had developed to the blastocyst stage (Table 2).

**Figure 4.**
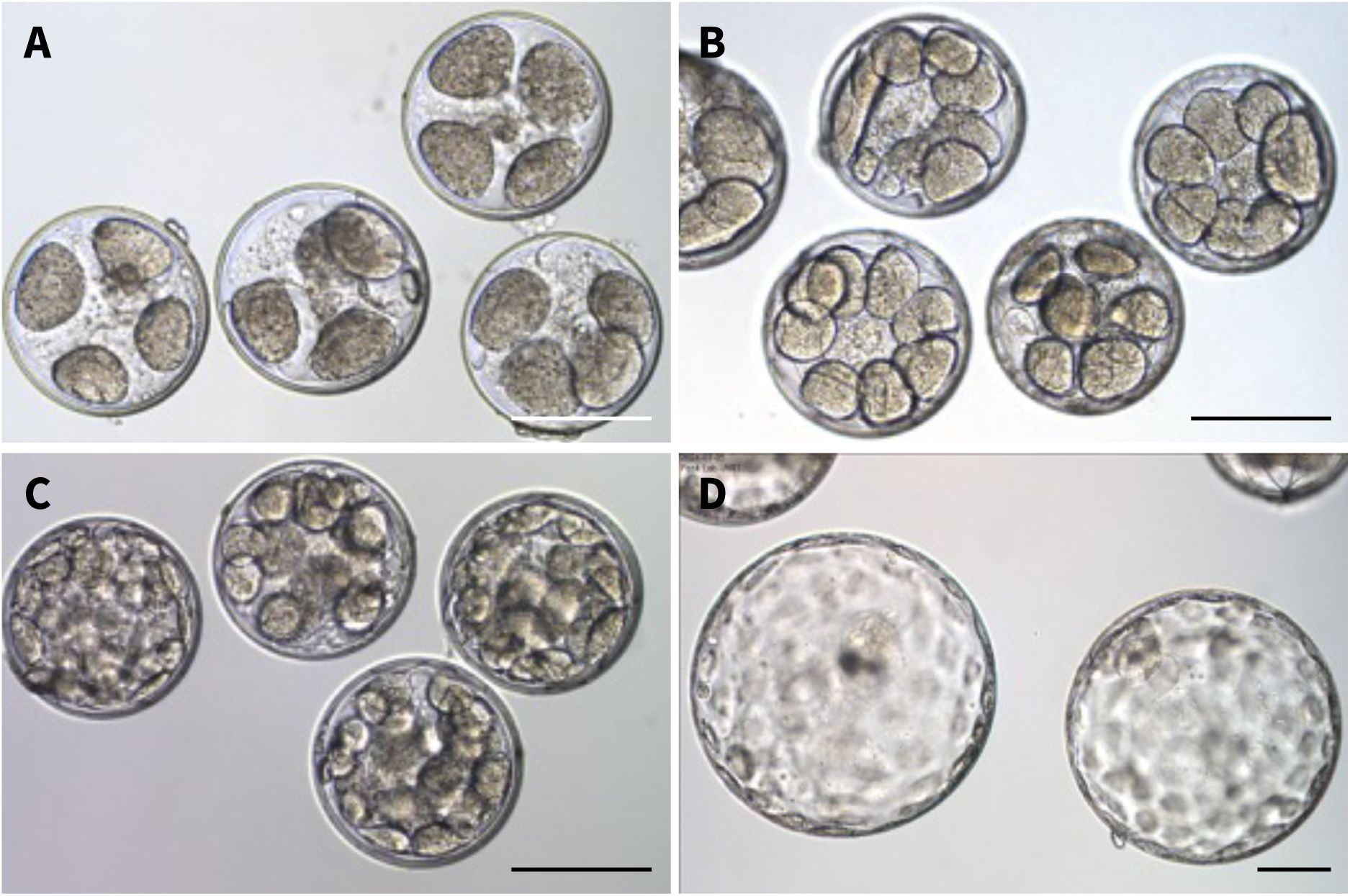
Prepubertal female dunnarts were primed with 1IU PMSG using the 3P6 treatment. Ovulation was induced by 1IU hCG 48 h post the third PMSG administration. The females were caged with stud male dunnarts for achieving fertilisation *in vivo*. 4-cell stage embryos (A) retrieved from uteri 48 h post-hCG injection. These developed *in vitro* in dKSOM1 medium to the 8-cell stage embryos (B), 16-cell or 32-cell stage embryos (C) 72 h post-hCG in dKSOM2 medium, and blastocyst stage embryos (D) 96 h post-hCG. Bars = 200 µm.

**Table 2.** Development capacity of dunnart embryos fer.lised in vivo.

| Experiment no. | No. of female | No. of 4-cell embryos retrieved | No. (%) of embryos developed into |  |
| --- | --- | --- | --- | --- |
|  |  |  | 8-cell/16-cell<sup>a</sup> | Blastocysts<sup>b</sup> |
| 1 | 1 | 23 | 8/8 (100) | 4 (50.0) |
| 2 | 1 | 20 | 9/9 (100) | 7 (77.8) |
| Total | 2 | 43 | 17/17 (100) | 11 (64.7) |

<sup>a</sup> % of number of 4-cell embryos. <sup>b</sup> % of number of 4-cell embryos.

### Validation of oestrus by vaginal lavage cytology

Cytology of vaginal lavages collected on the hCG injection day was characterized by almost exclusive detection of polygonal-shaped squamous cornified epithelial cells (CEC) in all treatment groups, indicating ovulation occurred. (Fig. 5).

**Figure 5.**
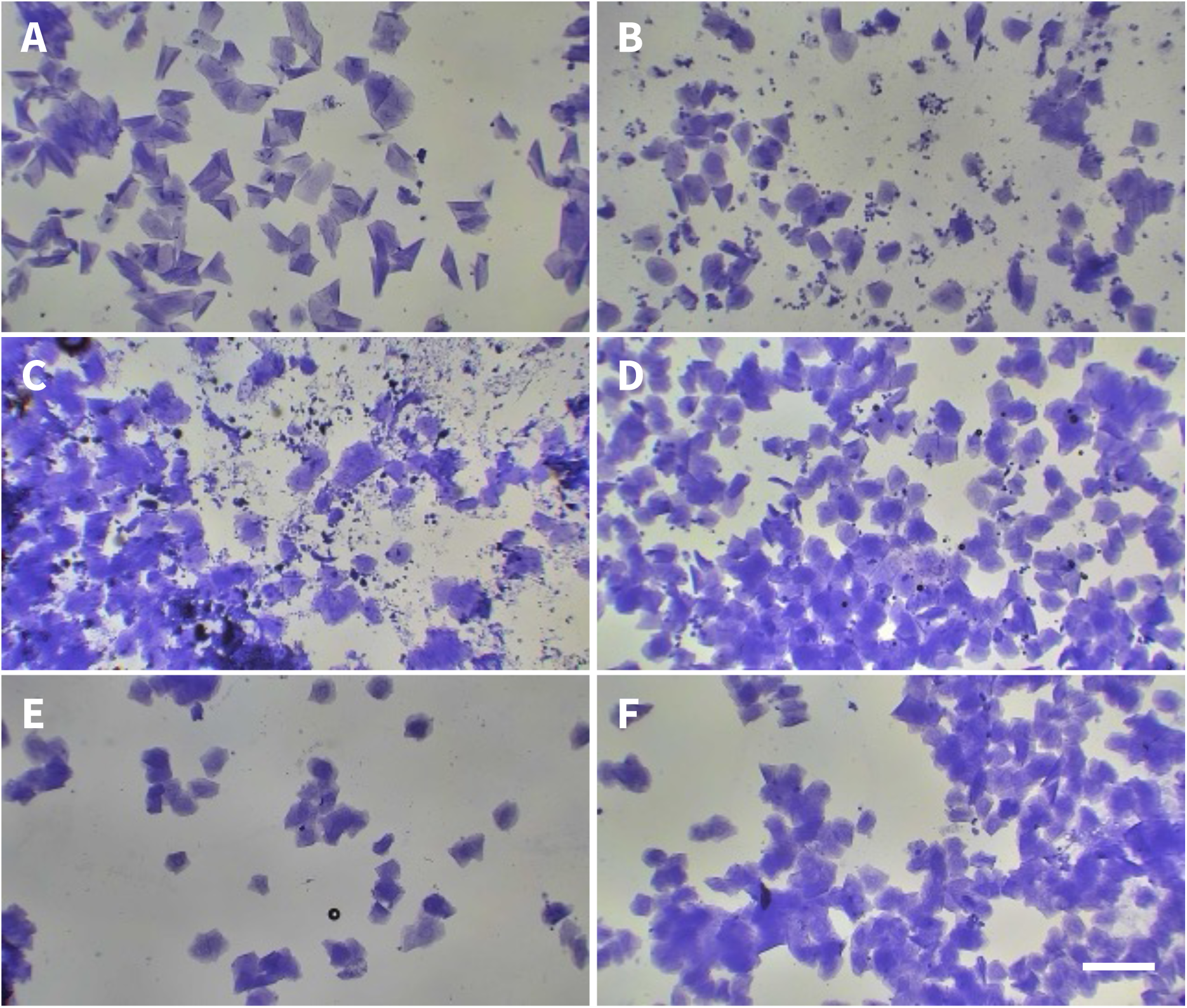
Cytological assessment of vaginal lavage samples from groups 1P4 (A), 2P4 (B), 1P6 (C), 3P6 (D), 1P8 (E) and 4P8 (F) following priming and collected at the time of hCG injection. The dominant cell type detected in each treatment group was squamous cornified epithelial cells (CEC), often in clumps. This indicted that females were coming into estrus and receptive to matings. Bar = 100 µm.

### Validation of ovulation by ovarian histology

Histological examination revealed that ovaries were dominated by corpora lutea (CL), indicating ovulation, in all treatment groups except the 1P4 group (Fig. 6B, C, E and F). While the ovaries from 1P4 and 2P4 treatment groups showed that the large antral follicles occupied significant areas of the tissue (Fig. 6A and D). The cumulus cells were separated from their companion oocytes after oocyte-cumulus complexes were extruded from the follicles (Fig. 6E). This explains why bare oocytes were retrieved from the ovulations.

**Figure 6.**
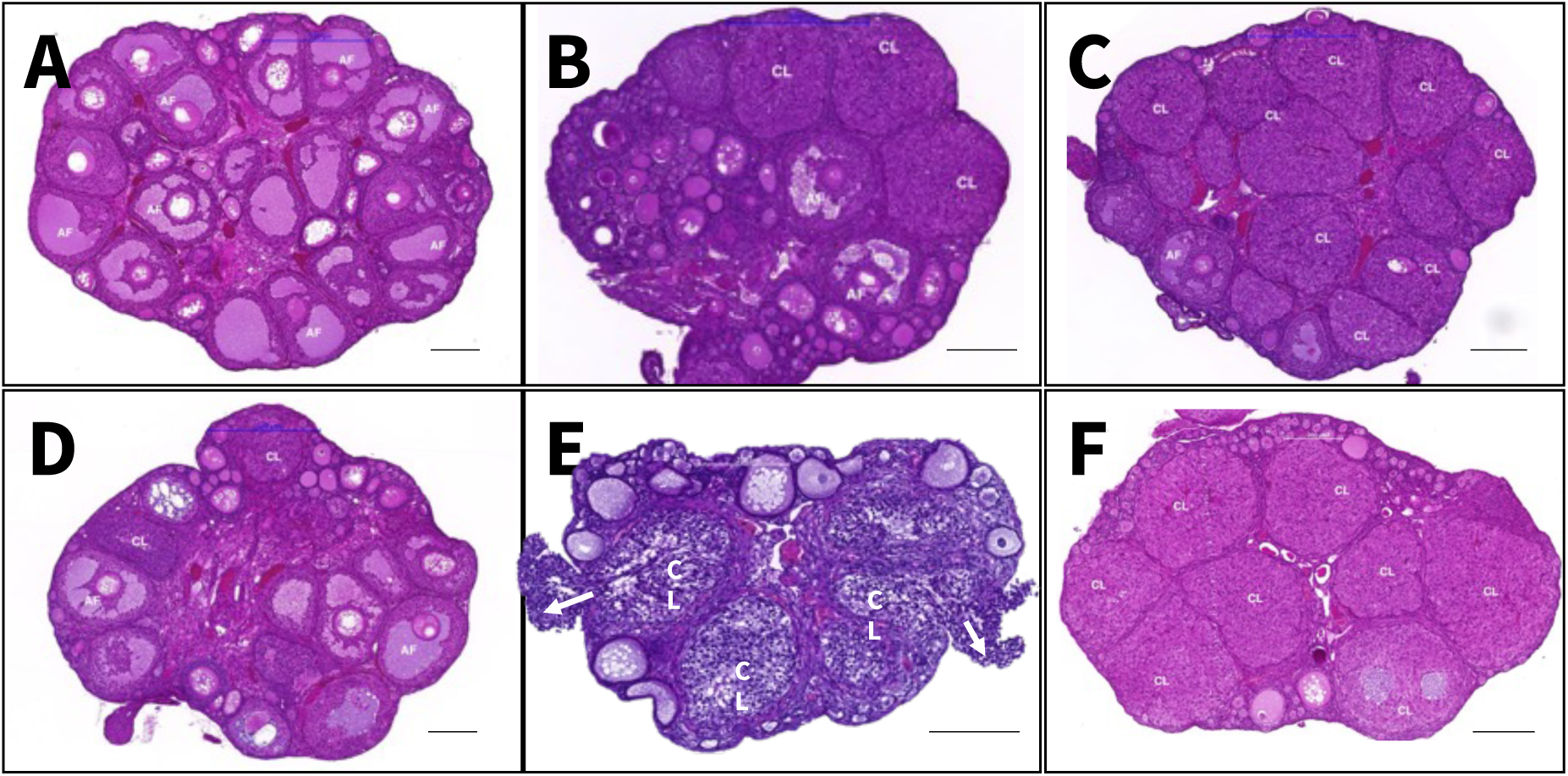
Representative histological sections of the ovaries from treatment group 1P4 (A), 1P6 (B), 1P8 (C), 2P4 (D), 3P6 (E) and 4P8 (F). Note the presence of large antral follicles (AF) and corpora lutea (CL) in the ovarian tissue. Cumulus cells (arrows in E) were extruded from the follicles during ovulation 4-4.5h post-hCG stimulation. Bars = 200 µm.

## Discussion

This study shows that mature and developmentally competent MII oocytes can be predictably retrieved from prepubertal dunnarts by a regimen of pregnant mare serum gonadotrophin (PMSG) and human chorionic gonadotrophin (hCG). The prepubertal female dunnarts aged 70-85 day old were primed with 1IU PMSG for 4-8 days to allow for follicle development from primordial follicles to preovulatory antral follicles. At the end of follicle development, the final oocyte maturation and ovulation were induced by administration of 1 IU hCG. In our present study, about 74% of prepubertal dunnarts responded to a 6-day PMSG stimulation and ovulated oocytes. More than 82% of the ovulated oocytes displayed normal morphology with a polar body in the thin perivitelline space and a clear deutoplast in the ooplasm. Moreover, the ovulated oocytes had the competence to be fertilised in vivo and had developed to 4-cell stage embryos by 48 h post-hCG administration, indicating the ovulated oocytes progressed meiosis to the MII stage. The majority of those 4-cell embryos (65%) went on to develop to the blastocyst stage in vitro after an additional 2-days of culture in dKSOM1 and dKSOM2 sequential medium.

The fat-tailed dunnart ovaries responded to all six treatment regiments. Four-day primed ovaries from both 1P4 and 2P4 groups were dominated with antral follicles with only a few corpora lutea. Six-day and eight-day primed ovaries contained all stages of growing follicles but were dominated by corpora lutea, indicating the occurrence of ovulation. Together, these data indicate that 4 days is not a sufficient priming window to enable reliable ovulations following hCG stimulation in the fat-tailed dunnart. The elevated appearance of polygonal cornified epithelial cells in vaginal lavage samples were observed in all groups, that indirectly reflected underlying circulating levels of ovarian steroids (oestradiol and progesterone) and gonadotrophins (FSH and LH) and indicated entry into the estrus phase of the reproductive cycle.

To determine the ideal stimulation regimen, we examined the optimal duration for follicle development to the ovulatory stage, stimulated by PMSG, and that produced the highest percentage of developmentally competent MII oocytes. The present study shows that ∼74% of the PMSG-primed prepubertal females ovulated with the highest rate of oocyte normality in the 3P6 group (3 injections over 6 days). Very few oocytes were recovered after 4 days stimulation while almost all prepubertal females primed for 8 days produced oocytes. However, most of the oocytes after 8 days stimulation were abnormal. This indicates that the follicular phase, in which primordial follicles developed to ovulatory stage follicles, is about 6 days in fat-tailed dunnarts. These results align with reports in a related marsupial species, the stripe-faced dunnart, *Sminthopsis macroura* [28, 40]. Indeed, degenerated oocytes and/or parthenotes, which probably were induced by endogenous LH, were retrieved from 8-day primed females (data not shown).

The animals ovulated 13 ± 11 (mean ± SD) oocytes per induced ovulation with the highest number of 35 oocytes retrieved after 6-day hormone priming. Moreover, the average number of normal oocytes was 11 with a standard derivation (SD) of 11. The large SD reflects the large variability seen in individual responses to the stimulation regimens. There are several explanations for this, the most likely of which is the developmental stage of the female stimulated. The prepubertal window chosen was day 70-85. This was chosen independent of animal weights or any other measures of maturity. Since this timpoint is on the cusp of their natural entry inot puberty, it is likely that at this early timepoint, some females were more responsive to the hormone stimulation protocols. While we could trial a different window, this might further confound data as some females might be too juvenile to enable stimulation or have already begun their normal cycling. This variations in the timing of puberty, oocyte number and normal oocyte rate is likely also counfounded by the polygenic background of our outbred dunnart colony. It is well established that genetic background is critically important to experiment outcomes, interpretation and reproducibility in rodents [41–43]. Thus, inbred strain mice have been most widely used in biomedical research and to mitigate the confounding effects of undefined variation [44, 45]. Moreover, the same concept of using inbred strains in research is applicable not only to mice and rats, but to a wide range of laboratory animal models, including rabbits, hamsters, guinea pigs and fish [45, 46]. It might be desirable to create inbred strains of the dunnarts by filial mating of brother and sister for multiple consecutive generations to achieve homozygosity to help further standardise protocols and decrease variability.

The competence and developmental potential of the oocytes produced in our study were assessed by in vivo fertilization and embryo culture to the blastocyst stage *in vitro*. The superovulated oocytes could be fertilized in vivo when the PMSG-primed females were mated with stud male dunnarts, and 4-cell stage embryos were collected 48 h post-hCG administration. The embryos could not develop further when cultured in commercial medium such as KSOM-aa (Merck), G1/G2 (Vitrolife) or BO-IVC (IVF Bioscience) (data not shown). Previous work had demonstrated that dunnart embryos had a much higher energy requirement than that of mouse, and glucose uptake was significantly greater at all stages of early embryo development in stripe-faced dunnarts than in mice [37]. Almost 65% of fat-tailed dunnart 4-cell embryos developed to the blastocyst stage when cultured in the high-glucose dKSOM1/dKSOM2 sequential medium [37] used in our study. Further investigation of embryo quality will examine total blastomere cell numbers, ratio between pluriblast and trophectoderm, and blastomeres apoptosis [47, 48].

In conclusion, the results reported here show that it is possible to induce ovulation in the prepubertal dunnarts using a PMSG priming regimen and obtain competent MII oocytes, which can be fertilized and develop to the blastocyst stage. Such an approach of superovulation establishes a protocol for the production of mature oocytes for use in future dunnart ARTs.

## Acknowledgements

The authors thank the staff at BioSciences 4 Animal Facility, the University of Melbourne, for the daily management of the dunnart colony. We acknowledge the Biological Optical Microscopy Platform at the University of Melbourne for the confocal microscopy. We acknowledge the Melbourne Histology Platform and Phenomics Australia Histopathology and Slide Scanning Service at the University of Melbourne for the assistance with tissue processing and imaging.

## Author contributions

JL–design of the project, data collection and analysis, and manuscript writing; NM–data collection and analysis, and manuscript editng; ELS–data analysis and manuscript editing; SO–design of the project and manuscript editing; AJP–design of the project, data analysis, and manuscript writing and editing.

## Data availability

Data are available on request.

---

**Supplemental Table S1.**
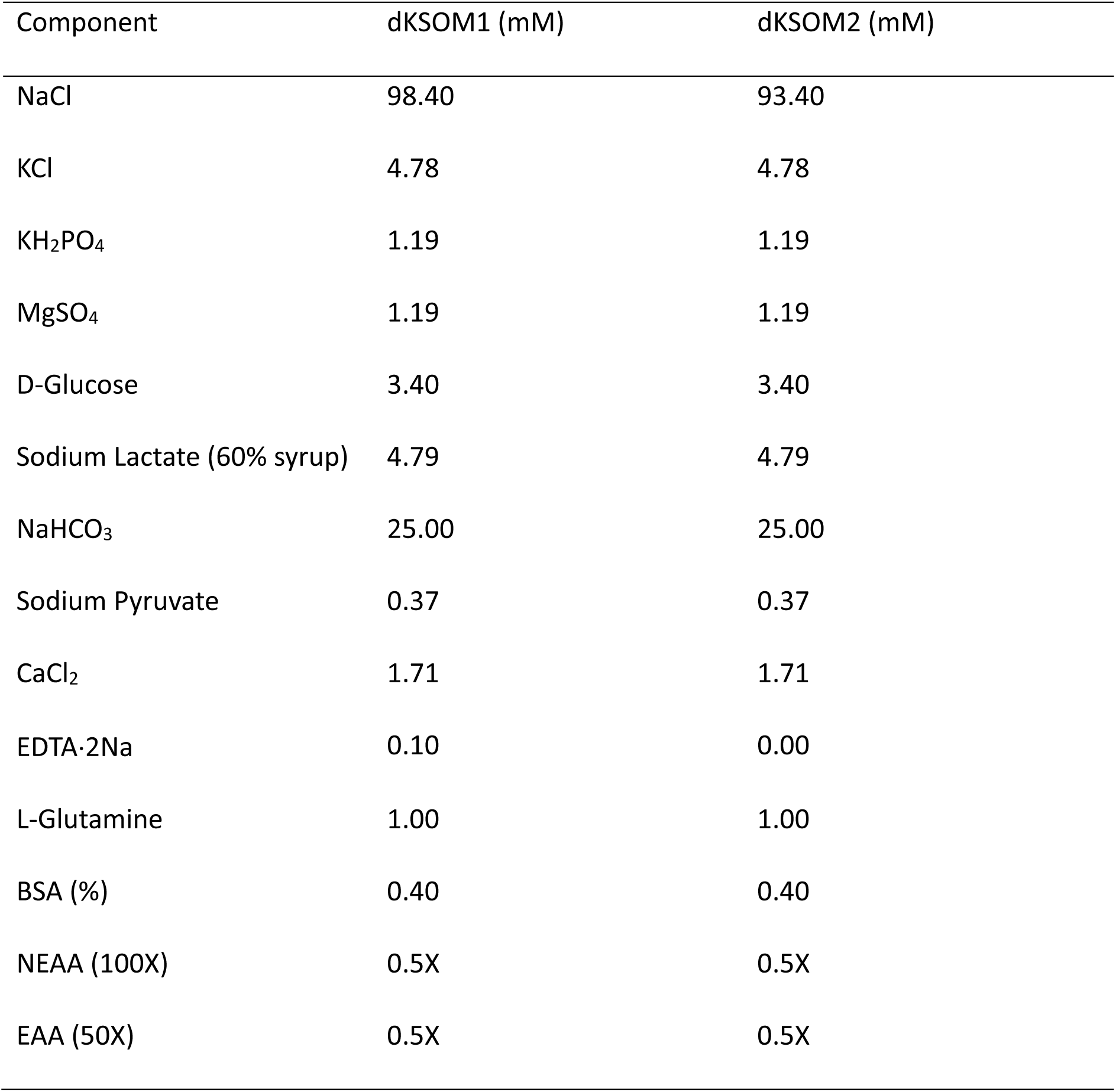
Composi.on of dKSOM1/dKSOM2 medium[37].

| Component | dKSOM1 (mM) | dKSOM2 (mM) |
| --- | --- | --- |
| NaCl | 98.40 | 93.40 |
| KCl | 4.78 | 4.78 |
| KH<sub>2</sub>PO<sub>4</sub> | 1.19 | 1.19 |
| MgSO<sub>4</sub> | 1.19 | 1.19 |
| D-Glucose | 3.40 | 3.40 |
| Sodium Lactate (60% syrup) | 4.79 | 4.79 |
| NaHCO<sub>3</sub> | 25.00 | 25.00 |
| Sodium Pyruvate | 0.37 | 0.37 |
| CaCl<sub>2</sub> | 1.71 | 1.71 |
| EDTA·2Na | 0.10 | 0.00 |
| L-Glutamine | 1.00 | 1.00 |
| BSA (%) | 0.40 | 0.40 |
| NEAA (100X) | 0.5X | 0.5X |
| EAA (50X) | 0.5X | 0.5X |

**Supplemental Table S2.**
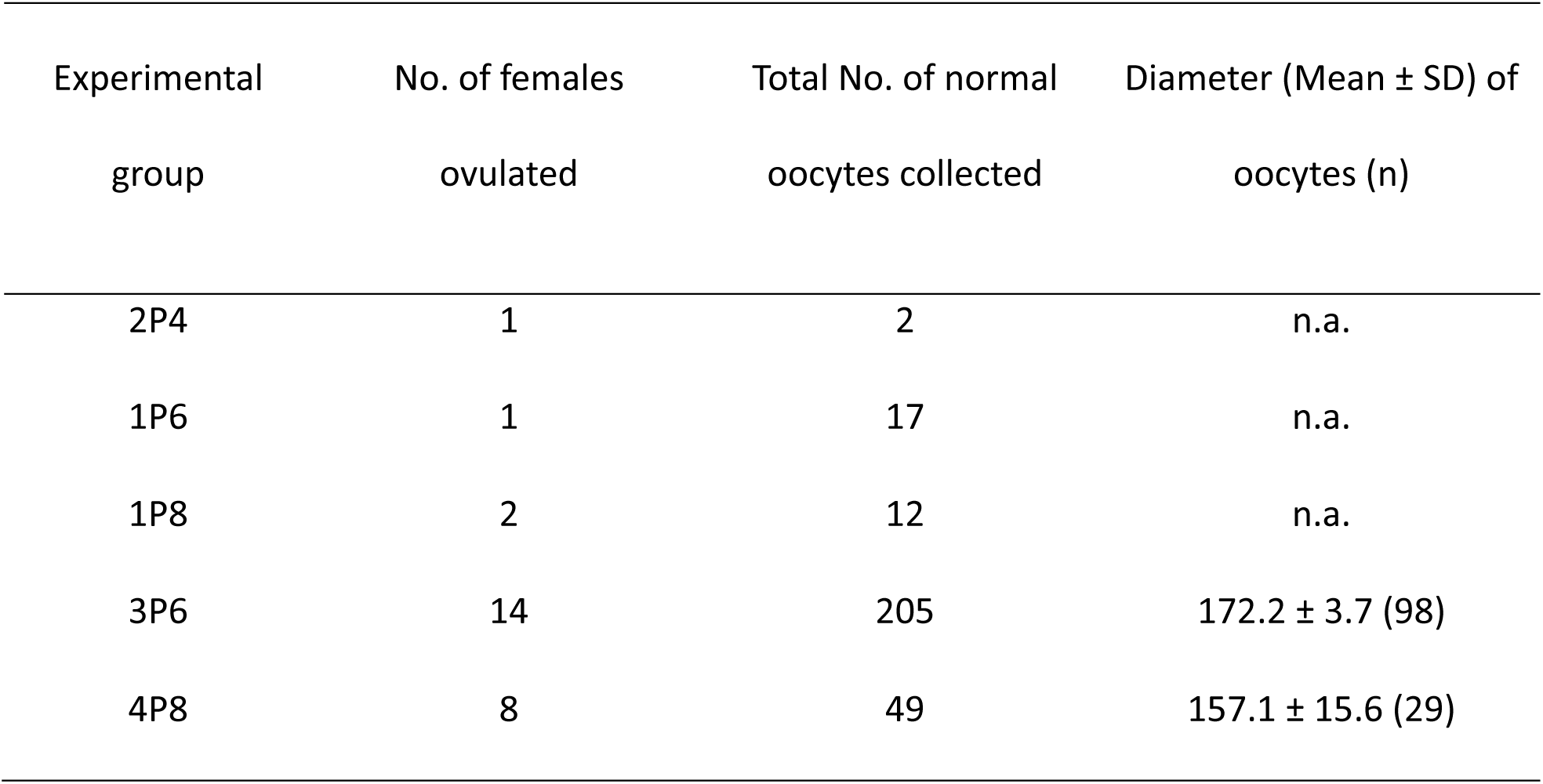
Total number of oocytes collected and diameter.

